# Feedforward amplification in recurrent networks underlies paradoxical neural coding

**DOI:** 10.1101/2023.08.04.552026

**Authors:** Kayvon Daie, Lorenzo Fontolan, Shaul Druckmann, Karel Svoboda

**Affiliations:** The Allen Institute for Neural Dynamics, Seattle, WA; Janelia Research Campus, Ashburn, VA; Turing Centre for Living Systems, Aix-Marseille University, INSERM, INMED U1249, Marseille, France; Stanford University, Stanford, CA

## Abstract

The activity of single neurons encodes behavioral variables, such as sensory stimuli (Hubel & Wiesel 1959) and behavioral choice (Britten et al. 1992; Guo et al. 2014), but their influence on behavior is often mysterious. We estimated the influence of a unit of neural activity on behavioral choice from recordings in anterior lateral motor cortex (ALM) in mice performing a memory-guided movement task (H. K. Inagaki et al. 2018). Choice selectivity grew as it flowed through a sequence of directions in activity space. Early directions carried little selectivity but were predicted to have a large behavioral influence, while late directions carried large selectivity and little behavioral influence. Consequently, estimated behavioral influence was only weakly correlated with choice selectivity; a large proportion of neurons selective for one choice were predicted to influence choice in the opposite direction. These results were consistent with models in which recurrent circuits produce feedforward amplification (Goldman 2009; Ganguli et al. 2008; Murphy & Miller 2009) so that small amplitude signals along early directions are amplified to produce low-dimensional choice selectivity along the late directions, and behavior. Targeted photostimulation experiments (Daie et al. 2021b) revealed that activity along the early directions triggered sequential activity along the later directions and caused predictable behavioral biases. These results demonstrate the existence of an amplifying feedforward dynamical motif in the motor cortex, explain paradoxical responses to perturbation experiments (Chettih & Harvey 2019; Daie et al. 2021b; Russell et al. 2019), and reveal behavioral relevance of small amplitude neural dynamics.

## Introduction

A central goal of neuroscience is to explain perception, decision-making and actions in terms of patterns of action potentials (Jazayeri & Afraz 2017). Neural recordings identify neurons with activity that is correlated with, or selective for, specific behavioral variables, such as tuning curves for visual stimuli (Hubel & Wiesel 1959) or behavioral choice (Britten et al. 1992). Large-scale recordings using silicon probes and cellular imaging have allowed trial-by-trial analysis of neural populations in the context of complex behaviors (H. Inagaki et al. 2019; Li et al. 2016). These analyses rely on dimensionality-reduction methods to identify activity subspaces that contain behavior-related selectivity. For example, in a memory-guided movement task, a single direction in activity space allows decoding of movement direction (choice) at the level of individual trials (H. K. Inagaki et al. 2018; Li et al. 2016). Similar analyses have been used to decode signals related to working memory (Machens et al. 2010; Murray et al. 2017), head direction (Chaudhuri et al. 2019), ego-centric position (Gardner et al. 2022), behavioral context (Mante et al. 2013), eye position (Daie et al. 2015), and task variables. In most cases the behavioral variables being encoded are low-dimensional and so are the corresponding features in the data (Gao et al. 2017). As a result, models linking neurophysiology and behavior are often low-dimensional (H. Inagaki et al. 2019; Li et al. 2016; Machens et al. 2010; Mante et al. 2013).

Dimensionality reduction methods by themselves do not reveal how neural activity at one time causes neural activity at later times and ultimately behavior. Moreover, it is not obvious that only subspaces containing the most variance in activity or selectivity have behavioral influence. Selectivity quantifies the correlation between neural activity and a behavioral variable and is determined by neuronal input. In contrast, behavioral influence is the behavioral impact per unit of neuronal activity, which in addition depends on connections to downstream target neurons. Selectivity can be estimated by recording activity during behavior. Behavioral influence is most directly measured by perturbing neural activity of specific neurons and observing the effects on downstream neurons and behavior. Efforts to measure behavioral influence of neural activity often suffer from a lack of specificity. Electrical microstimulation (Afraz et al. 2006; Salzman et al. 1990) and optogenetics (Guo et al. 2014; Pinto et al. 2019) have both been used to demonstrate a causal role for activity in specific brain areas, but these methods are unable to resolve which neurons were directly perturbed and which were recruited through synaptic connections.

Two-photon imaging of neural activity combined with two-photon optogenetics (Rickgauer et al. 2014) allows targeted perturbations of groups of individual neurons, providing for exploration of the relationship between selectivity and behavioral influence. Experiments in mice (Carrillo-Reid et al. 2019; Daie et al. 2021b; Jennings et al. 2019; Marshel et al. 2019; Russell et al. 2019) have shown that small groups 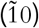 of neurons can bias behavior, but the relationship between neural selectivity and behavioral influence can be complex. For example, photostimulating V1 neurons tuned for one visual stimulus could evoke opposing percepts (Russell et al. 2019). Similarly, photostimulation of ALM neurons that were selective for one behavioral choice could bias the mouse to make the opposite choice (Daie et al. 2021b). Most analyses of targeted photostimulation experiments assume that subspaces containing selectivity and behavioral influence are aligned, implying that selectivity predicts behavioral influence. These experiments thus reveal an apparently paradoxical relationship between selectivity and behavioral influence, implying misalignment of the subspaces. In both V1 (Carrillo-Reid et al. 2019) and ALM (Daie et al. 2021b), behavioral biases were better explained by considering not just the targeted neurons, but also their effect on the surrounding neural population, highlighting the need to understand perturbation experiments in terms of interactions within neural circuits (Daie et al. 2021b; Finkelstein et al. 2021; Jazayeri & Afraz 2017; Li et al. 2016).

We use regression to identify activity subspaces that contain behaviorally influential activity in ALM of mice during a memory-guided, directional licking task. Neural dynamics evolved over multiple dimensions in activity space, culminating in choice-selective attractor dynamics (Finkelstein et al. 2021; H. Inagaki et al. 2019). Remarkably, early subspaces carried weak choice selectivity but had large behavioral influence, whereas late dimensions carried strong selectivity but weak behavioral influence. As a result, for individual neurons, behavioral influence was only weakly correlated with choice selectivity. These results were consistent with a neural circuit model (‘amplifying feedforward model’, AFF) in which weakly selective signals along early dimensions are amplified as they flow into later dimensions. We tested AFF model predictions by analyzing ALM dynamics and behavior in response to photostimulating small groups of neurons (Daie et al. 2021b).

### Identifying subspaces that influence behavior

Mice performed a memory-guided directional licking task (Fig. 1a) (H. K. Inagaki et al. 2018). Neurons in the anterior lateral motor cortex (ALM) exhibit choice-selective activity (preparatory activity) during the delay epoch, before the behavioral response (Fig. 1b). During late phases of the delay epoch, ALM preparatory activity is low-dimensional, with two dimensions accounting for more than 90% of selectivity (H. K. Inagaki et al. 2018) (Supp. Fig. 2). We define the coding direction (CD) as the direction in activity space that maximized selectivity at the end of the delay epoch (Fig. 1c). Activity projected to the CD (Fig. 1d) predicts behavioral choice on single trials under control conditions and under a variety of experimental manipulations (Finkelstein et al. 2021; H. Inagaki et al. 2019; Li et al. 2016). Choice selectivity and behavioral influence thus lie in the same subspace at the end of the delay epoch.

**Figure 1:**
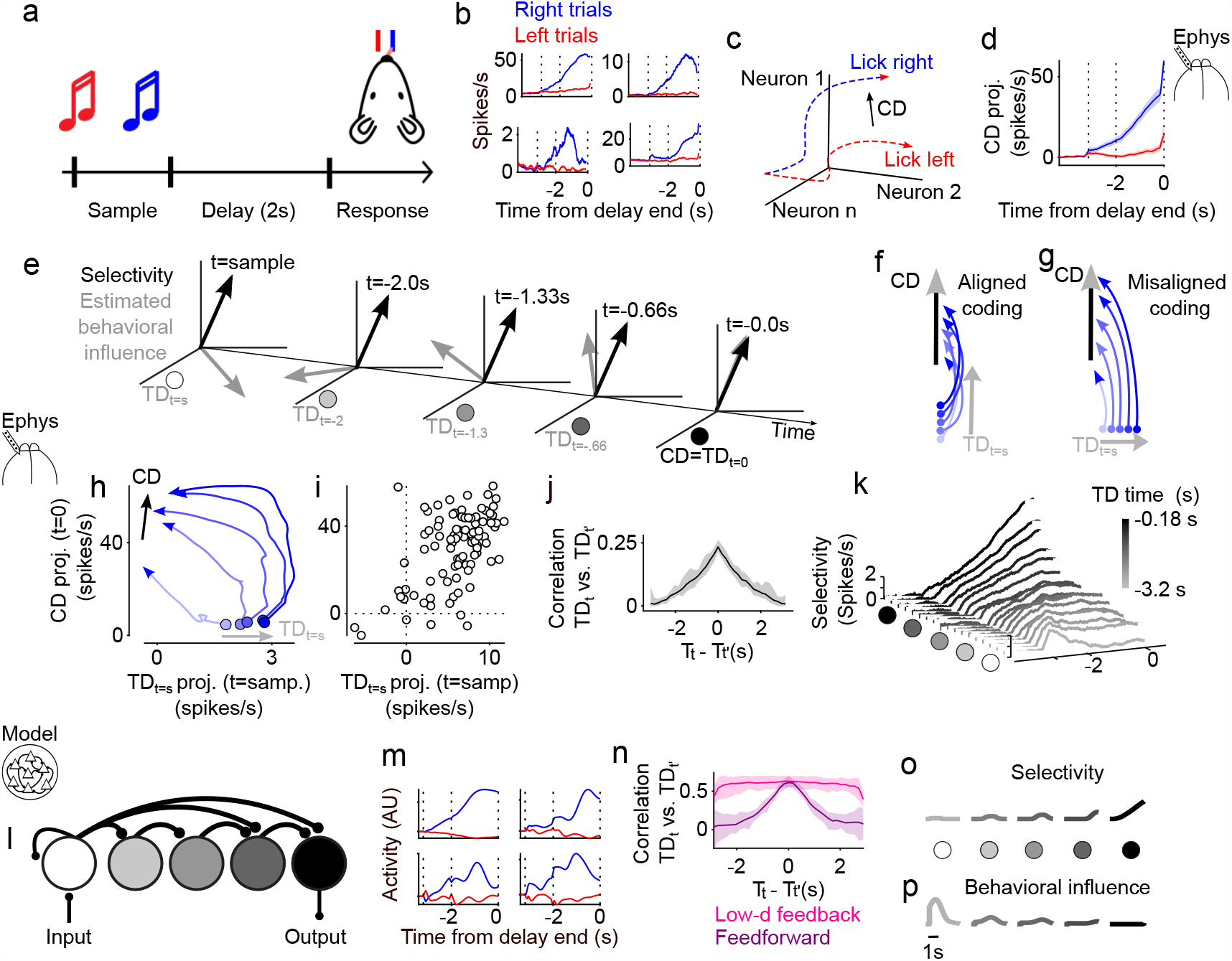
Multiple directions in activity space influencing behavior. **a**. Delayed directional licking task. Mice were instructed with auditory stimuli to lick left or right following a two second delay epoch. **b**. Spike rates of four simultaneously recorded single units on lick right (blue) and lick left (red) trials. These neurons are selective for rightward movements during the delay epoch. Dashed lines correspond to ticks in a. **c**. Schematic illustrating calculation of coding direction (CD). **d**. Trial-averaged activity projected along CD 19 recording sessions; 6 mice; 712 neurons; error shade, s.e.m. across sessions). **e**. Schematic illustrating estimation of the behavior influencing subspace. Behavioral influence (gray arrows) and selectivity (black arrows) are estimated separately at each time point. Panel depicts misaligned coding. Aligned coding would result in gray and black arrows overlapping at all time points. **f-g**. Schematic state-space trajectories in the *TD*_*t*=*sample*_ (*TD*_*t*=*s*_)-CD plane for aligned and misaligned coding models. Dark lines, average trajectories of trials with large-amplitude CD projections; light color lines, average trajectories of trials with small-amplitude CD projections. Circles, sample epoch; arrows, end of delay epoch. **h**. Same as f, g for electrophysiological recordings in ALM. During the sample epoch, activity along *TD*_*t*=*s*_ predicts activity in the late delay epoch along CD. **i**. Single trial activity along CD versus vs. *TD*_*t*=*s*_ for one behavioral session. **j**. Correlation between *TD*_*t*_ and *TD*_*t*_*′*, vs *t – t’* evaluated during sample and delay in 177.8 ms wide bins. All data were averaged across 19 recording sessions from 6 mice; Error shade, bootstrap 95% confidence interval. **k**. Choice selectivity along *TD*_*t*_. 18 *TD*_*t*_ were computed in 18 non-overlapping 177.8 ms wide bins spanning the sample and delay epochs. Selectivity is plotted along each *TD*_*t*_. Traces were offset by 0.2 spikes/s and the area under each curve was filled in with white shading to facilitate visualization. Traces were averaged across 19 experimental sessions from 6 mice. **l-n**. Amplifying feedforward (AFF) network model. **l**. Schematic of feedforward connectivity between *TD*_*t*_ (shades of gray) and the coding direction (CD; black). **m**. Activity of four right-selective example neurons from the AFF network. **n**. Correlation between *TD*_*t*_ and *TD*_*t*_*′*, vs *t – t’* in AFF model. Averages computed across 20 simulations.; Error shade, bootstrap 95% confidence interval. **o-p** Selectivity (o) (compare with k) and behavioral influence (p) along each activity direction in panel l. Electrophysiological data in b, d, h-k are from ref (H. Inagaki et al. 2019).

**Figure 2:**
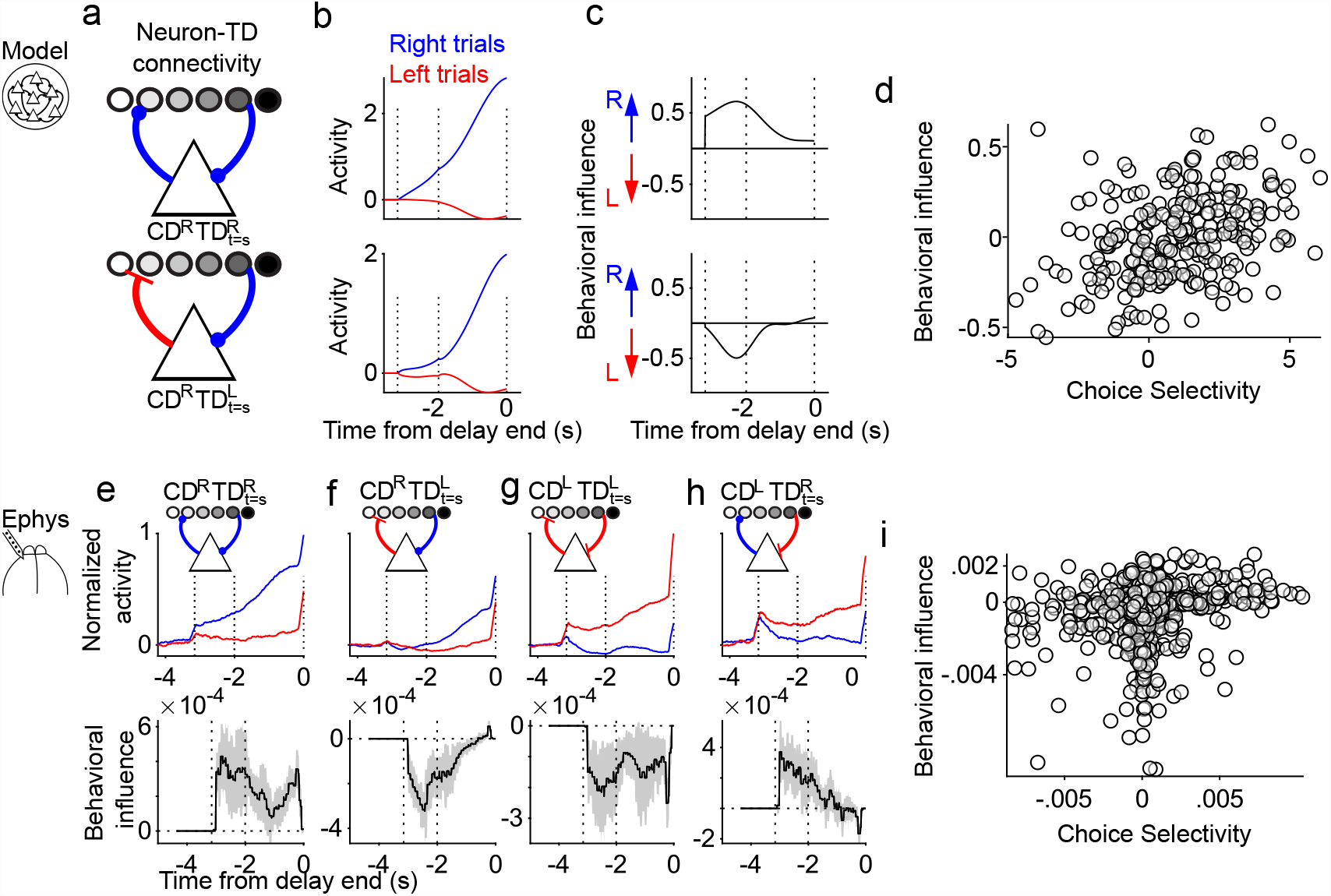
Misaligned coding in individual neurons. **a**. Connections between *TD*_*t*_ (circles) and neurons (triangles). Top: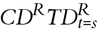 AFF model neuron is excited by late *TD*_*t*_ and excites early *TD*_*t*_. Bottom: 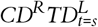 AFF model neuron is excited by late *TD*_*t*_ and inhibits early *TD*_*t*_. **b-c**. Activity on left (red) and right (blue) trials (b) and behavioral influence (c) of 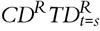 (top) and 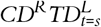(bottom) neurons. Behavioral influence of example neurons, bias at the network output following activation the example neuron. **d**. Time-averaged behavioral influence vs. selectivity of AFF model neurons. **e-i**. Behavioral influence estimated from ALM recordings. **g**. Time-averaged estimated behavioral influence vs. time-averaged selectivity for individual neurons (n=712 neurons). **e-h**. Average responses on left (red) and right (blue) trials (top) and behavioral influence (bottom) in groups of putative 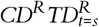 (neurons with right selectivity and rightward behavioral influence) (e), 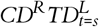 (f, n = 221 neurons), 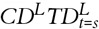 (g, n = 204 neurons), and 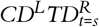 (h, *n* = 100 neurons). Error shade, bootstrap standard deviation.

We searched for directions in activity space at different time points during the behavior that may influence late-delay CD activity and thus behavioral choice. Using linear regression, we identified directions in activity space at time *t* that influence activity along the CD later in the trial (t = 0 s, end of delay epoch). We refer to these ‘transitional directions at time *t’* as *TD*_*t*_ (Fig. 1e). ‘Aligned coding’ implies CD parallel to *TD*_*t*_ for all *t* (Fig. 1f). ‘Misaligned coding’ (Fig. 1d-e) instead allows for *TD*_*t*_*′* with little overlap with the CD, and *TD*_*t*_*′* may vary over time (Fig. 1g). Misaligned coding implies that activity in directions other than CD predict behavior, especially earlier in the trial.

The earliest *TD*_*t*_ during the sample epoch, *TD*_*t*=*sample*_ (*TD*_*t*=*s*_) was nearly orthogonal to the CD, consistent with misaligned coding (Fig. 1h; range of distances, 77—89 degrees, 75% CI). Activity along *TD*_*t*=*s*_ during the sample epoch is predictive of activity along CD late in the delay epoch (Fig. 1i; avg. Pearson correlation for single sessions = 0.28; p = 0.002). Next, we computed *TD*_*t*_ in 18 non-overlapping time bins (177.8 ms). The resulting set of *TD*_*t*_ reside in an eight-dimensional volume in activity space (Supp. Fig. 1f; > 95% variance explained). To determine the time for the network to transition between *TD*_*t*_ we calculated the correlation between each *TD*_*t*_ with *TD*_*t′*_ at some other time point *t’* (Fig. 1j). Correlations dropped to 50 % of their peak within 810 ms (750 ms—820 ms, 75% CI across 18 *TD*_*t*_). Estimated behavioral influence is thus higher dimensional and more dynamic than choice selectivity, consistent with misaligned coding.

Selectivity for choice projected along each *TD*_*t*_ peaked at progressively later times for later *TD*_*t*_ (Fig. 1k). Additionally, selectivity grew in amplitude as the signal progressed along the *TD*_*t*_. Choice selectivity was 24.2 times greater for *TD*_*t*=0_ *≡CD* (18.2—32.5 times, 95% bootstrap CI) than for *TD*_*t*=*s*_ (Fig. 1k, Supp. Fig. 1e), implying amplification over time (Finkelstein et al. 2021; H. Inagaki et al. 2019; Li et al. 2016).

### Amplifying feedforward network model

Behaviorally influential activity sequentially traversed multiple directions in activity space. Similar sequential dynamics have been observed in feedforward network models with connections linking early directions to late directions (Ganguli et al. 2008; Goldman 2009). We constructed a feedforward network model (Fig. 1l) that produces dynamics matched to ALM activity (Methods). The network has three parameters: feedforward strength, feedback strength, and number of *TD*_*t*_. Feedforward connections set the strength of amplification: strong connections enable choice selective activity to grow as it propagates down the chain. Feedback connections from neurons contributing to each direction onto itself prolong the activity along that direction. Tuning the feedback parameter allows matching the timescales of transitions between *TD*_*t*_ in the data (Fig. 1j). Lastly, the number of *TD*_*t*_ in the model combines with the feedback parameter to set the duration of persistent activity in the network. This ‘amplifying feedforward’ (AFF) network produces dynamics that amplify signals as they propagate down the chain of *TD*_*t*_. Individual neurons exhibit selectivity that ramps up in time (Fig. 1m). Selectivity for choice is low-dimensional, as in ALM (Supp. Fig. 2b). The model exhibits *TD*_*t*_ transitions and dynamics similar to ALM (Fig. 1n; Fig. 1j) in an eight-dimensional subspace (Supp Fig. 1f).

Because signals are amplified as they progress down the chain, the early dimensions have a larger influence on behavioral output (per spike) than the late dimensions (Fig. 1o-p). To quantify behavioral influence, we perturbed activity along each direction in the model and calculated the response along the CD. Early *TD*_*t*_ had the largest influence on CD activity, despite having the weakest choice selectivity (Fig. 1o-p). The AFF network implements misaligned coding by decoupling behavioral influence and choice selectivity into distinct subspaces.

We also observed misaligned coding in recurrent neural networks (RNN) that were trained to produce ALM-like activity (Supp. Fig. 4). As with the AFF network, the RNN had low-dimensional selectivity by conventional measures, but showed multiple activity directions with significant behavioral influence which carried little variance in activity and little choice selectivity. The circuit mechanism underlying misaligned coding in the RNN was an interplay between amplifying feedforward connections and non-linear attractor dynamics along the CD (Finkelstein et al. 2021; H. Inagaki et al. 2019; Li et al. 2016). The AFF network (Fig. 1l) is a simple implementation of an ‘AFF dynamical motif’ that can be expressed by a variety of neural circuits.

### Misaligned coding in individual neurons

The AFF network has high behavioral influence along early *TD*_*t*_, including *TD*_*t*=*s*_, (Fig. 1p) and behavior-related selectivity along late *TD*_*t*_, including the CD (Fig. 1o). This misaligned coding causes apparently paradoxical coding at the level of individual neurons. We consider a hybrid view of connectivity in the AFF circuit, focusing on the interplay between individual neurons and *TD*_*t*_ (Fig. 2a). In this model, the selectivity of each neuron is determined by input from late *TD*_*t*_ (where choice selectivity is strongest), whereas their behavioral influence is determined by how strongly they project back to early *TD*_*t*_ (where behavioral influence is strongest) (Fig. 2b, c). Neurons with identical selectivity thus can exert opposing influence on behavior (Fig 2a-c). For example, neurons with right selectivity along the CD (*CD*^*R*^; Fig. 2a) can have behavioral influence that is to the right (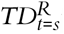 ; Fig. 2a, top) or left (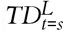; Fig. 2a, bottom). As a result, behavioral influence and choice selectivity of individual neurons are only weakly correlated (Fig. 2d).

We next analyzed neurons recorded in ALM for signatures of misaligned coding. The *i*^*th*^ element of *TD*_*t*_ is the estimated behavioral influence of neuron *i* at time point *t* (Fig. 2e-h). Similar to the AFF model, behavioral influence was largest early in the trial, whereas selectivity was largest late in the trial. Behavioral influence in ALM neurons is therefore only weakly correlated with selectivity (Fig. 2i, Pearson correlation = 0.09, p = 0.01). ALM contains neurons with behavioral influence in opposite directions from their choice selectivity (Fig. 2f,h), in addition to neurons that have aligned choice selectivity and behavioral influence (Fig. 2e,g).

### Directions with little choice selectivity can control behavior

ALM contains directions with small amplitude choice selectivity that are predicted by our analysis to have large behavioral influence. We next tested if the small amplitude signals along the behaviorally influential *TD*_*t*_ drive animal behavior. We analyzed calcium imaging experiments in ALM in which groups neurons were targeted for 2-photon photostimulation early during the delay epoch (Fig. 3a-b). Photostimulation produced biases in behavioral choice (Daie et al. 2021b). In a network with aligned coding, behavioral biases are expected to be correlated with the choice selectivity of the photostimulated neurons (i.e. projection to CD). For example, photostimulation of neurons with rightward CD selectivity should produce a rightward behavioral bias. In contrast, the behavioral bias was not correlated with the average CD selectivity of the photostimulated neurons (Fig. 3c, Pearson Correlation = 0.03, p = 0.35). Neurons with rightward CD selectivity caused both rightward and leftward biases, whereas neurons with leftward selectivity along the CD did not produce behavioral biases (Daie et al. 2021b) (Supp. Fig. 3).

**Figure 3:**
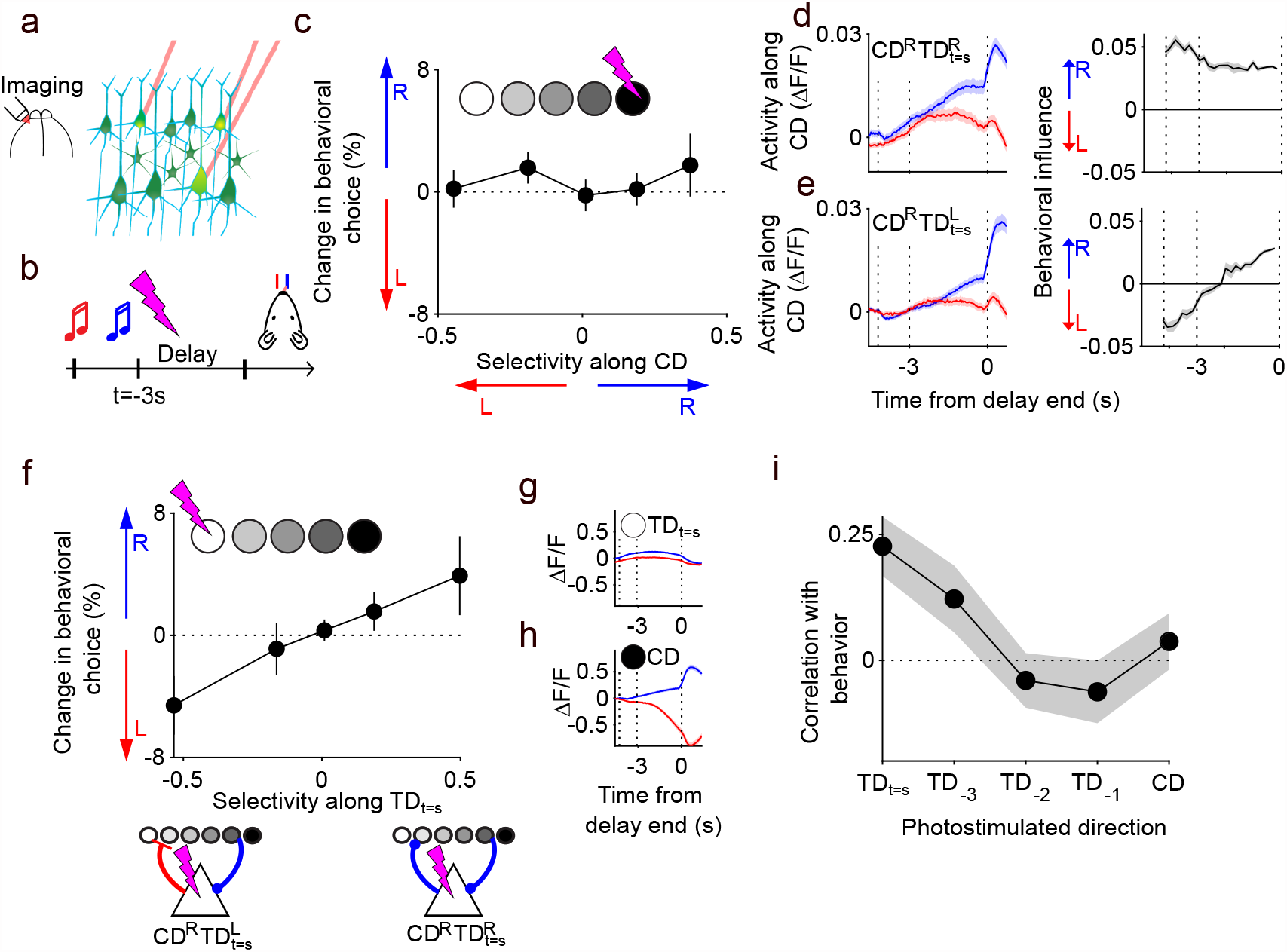
Directions with little choice selectivity control behavior. **a-b**. Schematic of targeted two-photon photostimulation. Brief (319 ms) photostimulation of 8 ALM neurons during the delay epoch (b). **c**. Change in behavioral choice vs. selectivity of directly photostimulated neurons along CD. Data were plotted in 5 equally spaced bins along the x-axis. Error bars,s.e.m across photostimulation groups. **d-e**. Trial averaged activity (left) and behavioral influence (right) of 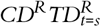 (d) and 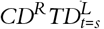 (e) neurons on right (blue) and left (red) non-photostimulation trials. Dashed lines, sample, delay and response epochs. Error shade, s.e.m across experimental sessions. **f**. Change in behavioral choice vs *TD*_*t*=*s*_ selectivity of the directly photostimulated *CD*^*R*^ neurons. Data were plotted in 5 equally spaced bins along the x-axis. Error bars, s.e.m across photostimulation groups. **g-h**. Average activity along *TD*_*t*=*s*_ (g) and CD (h). Blue, Right trials; Red, Left trials. Dashed lines, sample, delay and response epochs. **i**. Correlation between behavioral bias and selectivity (i.e., data in panels c & f) of the directly photostimulated neurons along *TD*_*t*_ and CD. Error shade standard deviation, bootstrap. Data averaged across 213 photostimulation groups from 84 behavioral sessions and 8 mice.

We tested if the estimated behavioral influence could predict which right selective neurons would produce rightward and leftward biases. As in the electrophysiological data, we found populations of neurons selective for right choice with rightward (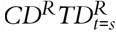, Fig. 3d) or leftward (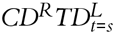, Fig. 3e) behavioral bias. 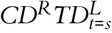 neurons represented 24% (avg, 18%—32%, 75% CI) and 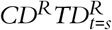 neurons represented 31% (avg, 20%—42%, 75% CI) of all imaged neurons. Behavioral bias was correlated with the *TD*_*t*=*s*_ selectivity of the directly photostimulated neurons (Fig. 3f, Pearson correlation = 0.22, p =0.0008). Given that *TD*_*t*=*s*_ carries only 4.8% (range, 4.7% — 4.9%, 75% CI, bootstrap) of the total sample and delay epoch choice selectivity (Fig. 3g,h), the large influence of *TD*_*t*=*s*_ on behavior is surprising, but similar to the AFF model. Photostimulation along later *TD*_*t*_ produced smaller behavioral biases (Fig. 3i). In the AFF model, the reduced behavioral influence of later *TD*_*t*_ is caused by amplification in the network; small inputs to *TD*_*t*=*s*_ are amplified, sequentially activating later *TD*_*t*_ ; inputs later in the chain are not amplified as strongly.

### Feedforward connectivity in ALM

In a network with the AFF dynamical motif, photostimulation along *TD*_*t*=*s*_ should trigger sequential activation of later *TD*_*t*_ in addition to producing behavioral biases. To test this prediction, we analyzed ALM responses following photostimulation along *TD*_*t*=*s*_. We identified photostimuli that activated 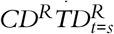 analyzed the responses of non-photostimulated neurons. Photostimulation of 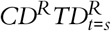 neurons (Fig. 4a-c) and neurons caused an increase in activity along *TD*_*t*=–1*s*_ in the non-photostimulated neurons that peaked at 1.2 seconds after photostimulation (range, 0.7 s–2.0 s, 75% CI; Fig. 4b). Photostimulation of 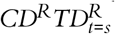 neurons also caused large amplitude ramping activity along the CD of the non-photostimulated neurons that peaked 1.9 seconds after the photostimulation (range, 1.0—2.2s, 75% CI; Fig. 4c). These photostimulation-evoked network dynamics were similar to the network activity observed during behavior (Fig. 1k). In contrast, photostimulation of 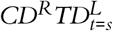 neurons (Fig.4d) produced weak inhibition along the *TD*_*t*_ and only a small change along the CD of the non-photostimulated neurons (Fig. 4e,f).

**Figure 4:**
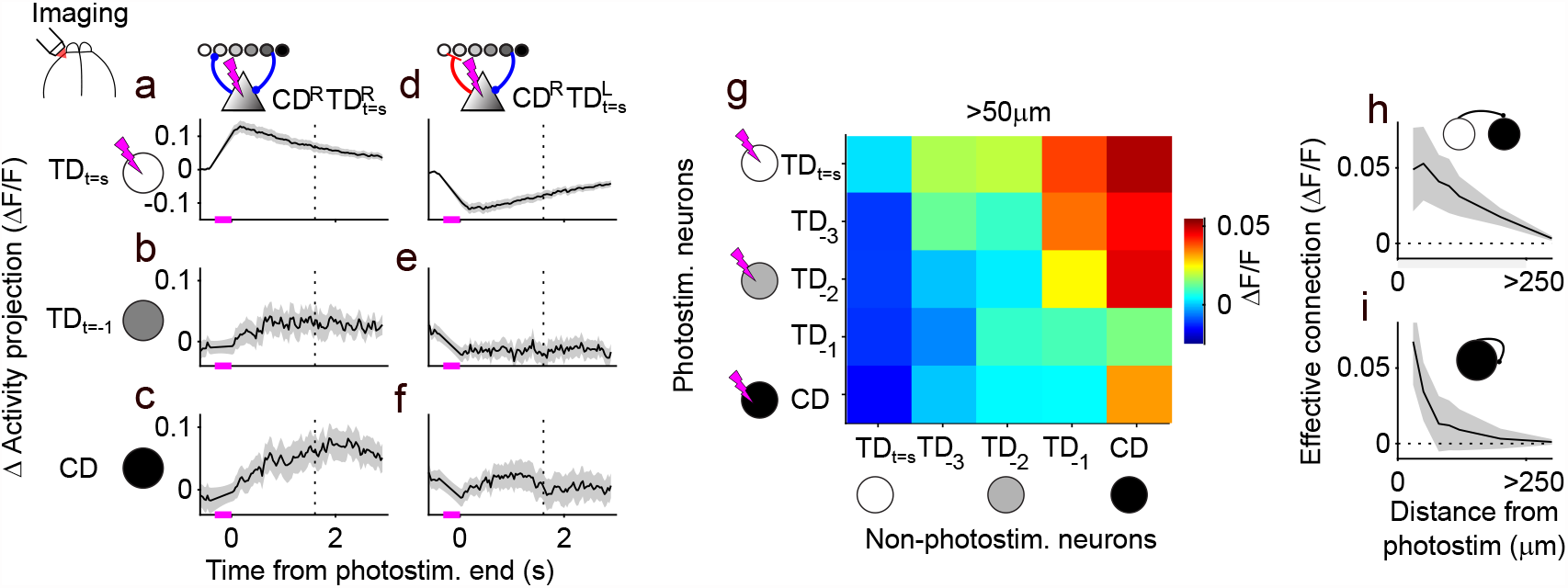
Feedforward connectivity in ALM. **a-f**. Δ Activity (difference in activity between photostim. And control trials) along *TD*_*t*=*s*_ following photostimulation of 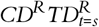(a)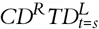(d)neurons. Activity projections for neurons located >50 fromphotostim. target along *TD*_*t*=–1*s*_ (b,e) and CD (c,f). Dashed line, end of delay epoch; magenta line, time of photostim.; Dashed line, go-cue. Positive values, lick right selective; negative values, lick left selective. Positive and negative activity projections along *TD*_*t*_ denote activation of neurons with right and left selectivity along *TD*_*t*_, respectively. **g**. Causal connection strength from *TD*_*t*_ onto *TD*_*t*_*′* for neurons >50 m from a target. **h**,**i**. Distance dependence of *TD*_*t*=*s*_ -CD (h) and CD-CD (i) causal connection strength. Error shade, s.e.m. from 40 photostimulation groups (eight neurons per group, 319 ms).

Changes in CD activity following photostimulation of *TD*_*t*=*s*_ demonstrate causal connections from *TD*_*t*=*s*_ to the CD. We estimated the strength of this connection by calculating the difference in the CD activity following photostimulation of 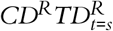 vs. 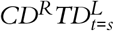 neurons (i.e. subtracting the traces in Figure 4c and f). We calculated causal connections between *TD*_*t*=*s*_, *TD*_*t*=–3_, *TD*_*t*=–2_, *TD*_*t*=–1_, and CD. Feedforward connections (early *TD*_*t*_ to late *TD*_*t*_) were stronger than feedback connections (late *TD*_*t*_ to early *TD*_*t*_) (Pearson correlation = -0.89, p = 3.1×10-9) (Fig. 4g-h). Additionally, the network has causal connections from the CD onto itself (Fig. 4i; Supp. Fig. 1). CD to CD feedback was prominent in local connections (30-50) (Fig. 4i), whereas longer range (50-300 μm) connections were primarily feedforward (Fig. 4h). These results reveal that ALM circuitry contains an AFF motif with spatial segregation of feedback and feedforward connections.

### Recovery of activity and behavior following large-scale photoinhibition

ALM dynamics are robust to large-scale photoinhibition of network activity via attractor dynamics (H. Inagaki et al. 2019) (Fig. 5a). In one-dimensional attractor models (Li et al. 2016), elimination of selectivity along CD by photoinhibition eliminates memory and scrambles trajectories. We calculated the correlation between CD activity and behavioral choice before, during and after photoinhibition (Fig. 5b; data from (H. Inagaki et al. 2019)). CD dynamics became uncorrelated with behavior during photoinhibition. CD activity before photoinhibition was correlated with behavior (Fig. 5b); this suggests that although CD choice selectivity was transiently eliminated during photoinhibition, choice information must have been preserved in the network.

**Figure 5:**
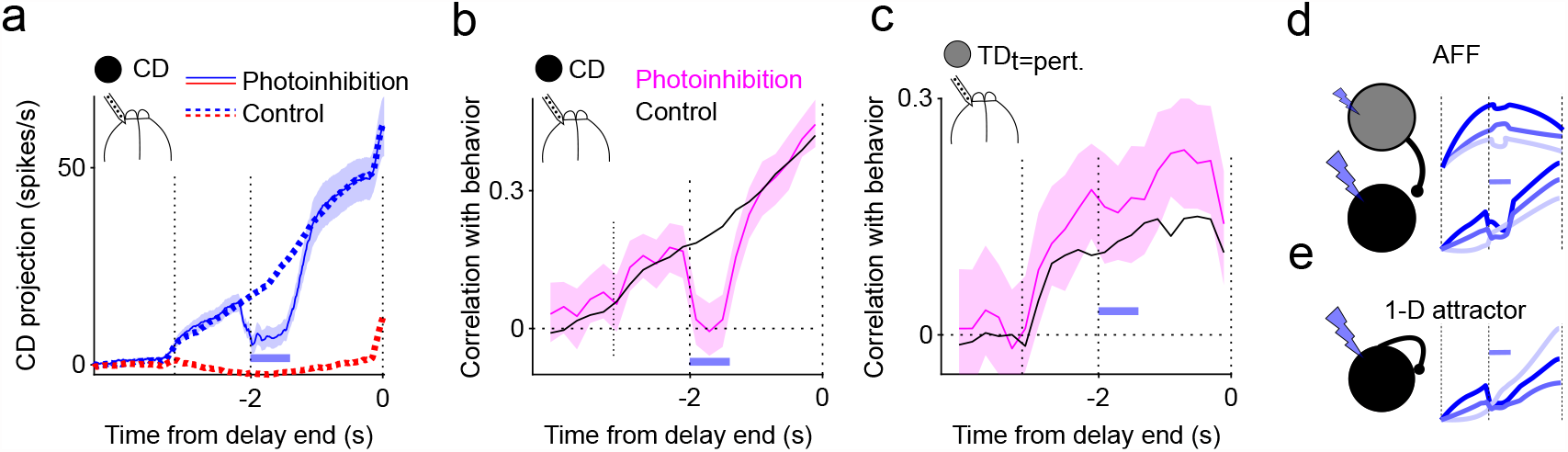
Recovery of activity and behavior after large-scale photonihibition. **a**. Activity along CD on control (dashed), lick right (blue), lick left (red) trials and lick right photoinhibition trials (solid) **b-c**. Correlation between activity projections along CD (b) and *TD*_*t*=*pert*._ (c) during control (black) and photoinhibition (magenta) trials. Blue bars, time of photoinhibition. **d-e**. Schematic single-trial dynamics following photoinhibition in an AFF (d) and one-dimensional attractor (e) model.

How does activity along CD and behavior recover after photoinhibition? In a higher-dimensional system, memory could remain intact during photoinihbition orthogonal to CD to aid correct recovery of activity along the CD and thus robustness. We identified a subspace in activity space that was not impacted by photoinhibition (*TD*_*t*=*pert*._; Methods). Activity along *TD*_*t*=*pert*._ during photoinhibition can be used to predict the choice equally on photoinhibition and control trials (Fig. 5c, bottom, p>0.92). These results are consistent with the presence of an AFF dynamical motif (Fig. 5d; Supp. Fig. 5), but not the simplest one-dimensional attractor model (Fig. 5e), suggesting that robustness of ALM dynamics could be facilitated by small amplitude signals that are causally linked to behavior, but unaffected by the perturbation.

## Discussion

In a memory-guided movement task, ALM dynamics are low-dimensional, with choice selectivity mostly contained along a single coding direction (CD) (H. K. Inagaki et al. 2018; Li et al. 2016). Here we identified a behaviorally influential subspace with higher dimensionality. Within this subspace, task-relevant information sequentially flowed through multiple directions into the CD. Previous studies established that activity along the CD during the delay epoch determines behavioral choice (Li et al. 2016). We found that activity in different directions in activity space was causally related to choice, as early as the sample epoch, seconds before the behavioral report. These results are surprising because these dimensions can contain little choice selectivity. The influence of these directions on behavior was explained by a model (AFF) with amplifying feedforward connections. Targeted photostimulation experiments revealed that ALM contains amplifying feedforward connectivity, as predicted by the AFF model.

Dimensionality reduction methods identify directions in neural activity space that explain high variance and are, by construction, correlated with behavior (Kobak et al. 2016; Mante et al. 2013). Activity is often reported to be low-dimensional, and the data are modeled with low-dimensional neural networks (Machens et al. 2010; Mante et al. 2013). This approach assumes that large amplitude signals exert the largest causal influence on behavior. Our results instead reveal that directions in activity space with relatively little variance in activity carry information that has a strong influence on behavior. We explain these results using a “dynamical systems perspective” (Shenoy et al. 2013), which shifts the focus from selectivity of individual neurons toward a focus on the influence that each neuron (or group of neurons) exerts on the rest of the network. We verified predictions from a dynamical systems model using data from targeted photostimulation experiments (Figs. 3-4).

The AFF network is an example of a broader class of “non-normal” networks. Non-normal networks have been shown to optimize signal-to-noise ratio during short-term memory (Ganguli et al. 2008). Variations on the AFF motif have also been used to model amplification of stimuli in visual cortex (Murphy & Miller 2009), planning and execution of movements (Hennequin et al. 2014), high-fidelity information transfer between neural populations (Baggio et al. 2020), and to enhance expressivity in recurrent neural networks for machine learning applications (Kerg et al. 2019).

ALM exhibits attractor dynamics that are robust to large-scale perturbations (Finkelstein et al. 2021; H. Inagaki et al. 2019). This robustness is based on two stable network states along the CD that are maintained by feedback. Previous attractor models require external ramping signals with weak selectivity to produce ALM activity patterns. This is an example of misaligned coding because the non-selective ramping direction influenced CD activity through non-linear amplification (Finkelstein et al. 2021; H. Inagaki et al. 2019). In addition, CD attractor dynamics are influenced by the AFF motif. The AFF motif can itself produce dynamics resembling a discrete attractor. Because of sequential amplification, the AFF is only sensitive to perturbations along *TD*_*t*=*s*_. Therefore, dynamics along the later *TD*_*t*_ are robust to perturbations. Combining the AFF motif with a discrete attractor circuit could potentially enhance stability of the dynamics with respect to perturbations. Future work involving perturbations to larger groups of neurons along the CD and neurons in deeper layers of ALM will be needed to disentangle the relative contributions of feedback amplification and amplification via the AFF motif.

Targeted photostimulation experiments have uncovered signatures of misaligned coding in visual cortex (Chettih & Harvey 2019; Russell et al. 2019). We propose that similar dynamical motifs might underlie these apparently paradoxical results. In one study, neurons in V1 were found to surprisingly suppress their co-tuned neighbors (Chettih & Harvey 2019). Relatedly, it was found that photostimulation of neurons tuned for a particular visual stimulus would sometimes enhance stimulus detection (as expected), but other populations of similarly tuned neurons would suppress stimulus detection (Russell et al. 2019). These results are consistent with a two-dimensional feed-forward network (Murphy & Miller 2009) in which stimulus tuning is determined by inputs from the late direction (i.e. CD), and behavioral influence is determined by connections onto the early direction (i.e. *TD*_*t*=*s*_). Consistent with this idea, behavioral biases caused by V1 photostimulation could best be explained by considering the influence of target neurons on the surrounding population (Carrillo-Reid et al. 2019).

Advances in the scale and precision of neurophysiological recordings (Campbell et al. 1991; Jun et al. 2017; Sofroniew et al. 2016) have motivated new theories (Chung & Abbott 2021; Gallego et al. 2017; Jazayeri & Afraz 2017) and methods (Churchland et al. 2012; Kaufman et al. 2014; Kobak et al. 2016; Li et al. 2016; Low et al. 2018; Machens et al. 2010) for decoding and interpreting population activity. A variety of methods have been used to uncover potentially useful information in small-amplitude correlations between pairs or groups of neurons (Nieh et al. 2021; Pandarinath et al. 2018; Schneidman et al. 2006). Here, we provide evidence that the directions carrying these correlations exert a strong influence on behavior despite accounting for only a small fraction of selectivity. Our findings highlight the potential insights to be gained from analyzing simultaneous recordings of large neuronal ensembles from a dynamical systems perspective.

## Methods

### Electrophysiological recordings

Electrophysiological data (Figures 1, 2 & 5) are from (H. K. Inagaki et al. 2019; H. Inagaki et al. 2019; H. K. Inagaki et al. 2018). Silicon probe recordings in the left hemisphere of ALM were performed in 19 behavioral sessions across 6 mice with an average of 37 neurons (range, 23—57). Spike rates of the *i*^*th*^ neuron during the *tr*^*th*^ trial at time *t*, 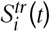, were computed using a 200 ms moving window. Time *t* was calculated relative to the end of the delay epoch. Choice selectivity of neuron *i* at time *t* was calculated as the average spike rate on correct right trials minus the average spike rate on correct left trials

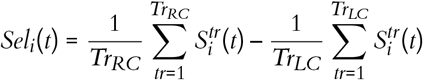

where *Tr*_*RC*_ is the number of correct lick right trials and *Tr*_*LC*_ is the number of correct lick left trials.

### Identifying subspaces that influence behavior

We developed a regression-based analysis to identify the transitional directions at time *t* (*TD*_*t*_) with the largest behavioral influence at each time point, *t* (Figs. 1-5). First, we calculated the Coding Direction (CD) as the average choice selectivity at the end of the delay epoch (i.e. *t* = 0 s) (*CD ≡ TD*_*t*=0*s*_). The expression for the *i*^*th*^ entry of the CD vector, corresponding to the weight of the *i*^*th*^ neuron reads

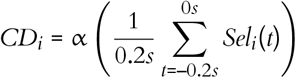

The normalization factor α was included to ensure that CD has unit norm. Activity projected along the CD at the end of the delay was given by

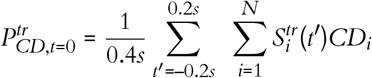

where *N* is the number of neurons. We used linear regression to find the neuronal subspace active at time *t* that best predicted the CD projections at the end of the delay epoch across trials, such that

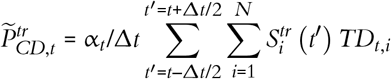

where 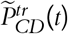 is the estimate of late-delay CD activity using activity in the time bin from *t* – Δ*t* to *t* + Δ*t*. The scalar α_*t*_was calculated to ensure that *TD*_*t*_ had unit norm. *TD*_*t*=*s*_ was defined as the first *TD*_*t*_ evaluated during the sample epoch (–3.15*s* < *t* < –2*s*, electrophysiology; –4.25*s* < *t* < –3*s*, 2-photon imaging). *TD*_*t*_ was calculated using the pseudoinverse (built-in MATLAB function *pinv*). Regression was done separately for left and right trials. All analyses in the paper focus on *TD*_*t*_ determined on right trials because activity modulations during right trials are much larger than on left trials in the left hemisphere of ALM. The estimated behavioral influence (Fig. 2e-i) at time *t* was the unnormalized *TD*_*t*_ (i.e. 1/α_*t*_*TD*_*t*_). Plots of behavioral influence vs. time (Fig. 2e-h) were computed in 80 ms wide bins. Trials were split into non-overlapping sets. One half of trials was used for assigning neurons as having rightward or leftward behavioral influence (training) and the other half was used for plotting (testing). Data were randomly split into testing and training halves ten times. Error shades represented the standard deviation across these ten repetitions.

To display 2D activity projections (Fig. 1h), we first orthogonalized *TD*_*t*=*s*_ and CD and then computed the projection at each time point for each trial. Trials were then sorted according to the amplitude of 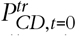 and then averaged in five bins each containing equal numbers of trials. For example, trajectories from all trials whose 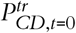 was in the 80^th^ percentile were averaged together (Fig. 1h; darkest line). To test whether activity along the TD_t_ at time point *t* could be used to predict late-delay CD activity we plotted 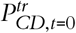 vs. 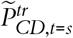 and calculated the Pearson correlation coefficient (Fig. 1i). In all analyses, non-overlapping halves of trials were used for independently calculating vectors (*TD*_*t*=*s*_ and CD) and projections.

Next, we calculated TD_t_ in 18 equally spaced time bins (177.8 ms) between the start of the sample epoch and the end of the delay. This bin size was chosen to approximately match the time window used for computing spike rates (200 ms). To cross-validate our results, we split trials into two non-overlapping sets to obtain two independent calculations of TD_t_ at each time point. We calculated the correlation between these two estimates TD_t,half1_ and TD_t,half2_ at each time point (Fig. 1j). To estimate the dimensionality of the space containing all 18 TD_t_ vectors we used a singular value decomposition. We computed the SVD of the average 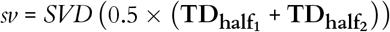 where 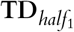 is a matrix with columns containing all 18 TD*t* computed from the first random half of trials. Next, we computed an SVD of the difference 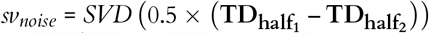 to estimate the contribution of trial-to-trial fluctuations to the dimensionality (Daie et al. 2015; Machens et al. 2010). The differences between *sv* and *sv*_*noise*_ were then used to calculate the dimensionality. The fraction of the full set of vectors *TD*_*t*_ contained along each SVD direction *k* was then given by 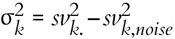. Finally, we calculated the normalized cumulative sum as 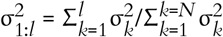. The dimensionality of *TD*_*t*_ was then defined as the first value of 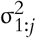 to be greater than 0.95. To estimate the dimensionality of the trial-averaged choice selectivity (Supp. Fig. 2), we repeated this analysis but instead computed the singular values of choice selectivity *Sel*_*i*_(*t*) at time point *t*. To verify that our regression analysis was capable of distinguishing between feedback and feedforward networks, we repeated all analyses on our AFF, FB only and FF only models (Supp. Fig. 1-2).

To verify that *TD*_*t*_ represented an accurate estimate of behavioral influence in network models, we perturbed each neuron in both an AFF network and an FB only network. The behavioral influence of neuron *i* was defined as the change in CD activity caused by the perturbation of neuron *i*. We then compared the behavioral influence of each neuron to the average of *TD*_*t*_ across all time points (i.e. 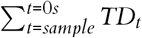; y-axis Supp. Fig. 1j-k).

### Analysis of photoinhibition experiments

Transient perturbations via bilateral photoinhibition of ALM were delivered on 25% of trials during the first 600 ms of the delay epoch (H. Inagaki et al. 2019) (Fig. 5). To identify the component of *TD*_*t*=*pert*._ that was unaffected by photoinhibition, we first calculated the difference in spike rate on photoinhibition and non-photoinhibition trials and used an SVD to find the first dimension that contained the largest changes in activity resulting from photoinhibition. Contributions from this direction were then subtracted from the population responses, and we calculated *TD*_*t*=*pert*._ using the regression approach described in the previous section. We then calculated activity projections along both *TD*_*t*=*pert*._ and *Sel*_*i*_(*t*) at each time point for both non-photoinhibition and photoinhibition trials. Next, we calculated the correlation between activity and behavioral choice along both *CD* (Fig. 5i) and *TD*_*t*=*pert*._ (Fig. 5j) throughout the trial. These analyses were then repeated for the AFF and one-dimensional attractor models (Supp. Fig. 5).

### Network models

Activity in model networks (Fig. 1, 2 & 5; Supp. Figs. 1-5) was governed by a system of equations

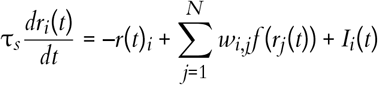

where *r*_*i*_(*t*) is the spike rate of neuron *i*, τ_*s*_ is the synaptic time constant, *w*_*i,j*_ is the strength of the connection from neuron *j* onto neuron *i*, and *I*_*i*_(*t*) is the strength of the external input onto neuron *i* and *f* (*x*) was the synaptic non-linearity. The input *I*(*t*) was set equal to one during the sample epoch and zero at all other times. Linear neuronal activations (i.e. *f* (*x*) = *x*) were used for all models except for the one-dimensional attractor network in Supp. Fig. 5 which had *f* (*x*) = 5 (tan *h*(*x*/10) + 1). To construct a network with an amplifying feedforward structure (Fig. 1l) we used the Schur decomposition (Strang 1993) such that

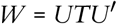

where *W* is the (*N×N*; N=150) connectivity matrix, *U* is the orthonormal set of *m* vectors corresponding to the Schur directions and *T* is an *m×m* upper triangular matrix whose entries represent the connections between the Schur directions. The number of vectors, *m*, determines the depth of the amplifying feedforward chain. We chose *m = 60* for the AFF network to replicate the finding that ALM *TD*_*t*_ spanned an eight-dimensional subspace (Supp. Fig. 1). For the feedback only network we used *m =* 150 and for the AFF network we used *m = 60*. For model networks, we observed a monotonic relationship between *m* and *TD*_*t*_ dimensionality (not shown). *U* was generated using a random initialization. The last column, *U*_*m*_, represented the output of the network and was chosen to be all-positive random values sampled from a uniform distribution, except in Supp. Fig. 5a,c,e in which all entries were 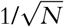 in order to maximize the overlap of the CD with the direction of photoinhibition. To ensure that the first Schur direction had equal numbers of neurons with positive and negative weights, the *i*^*th*^ entry of the first column of *U* was: *U*_*i*,1_ = *a*(*i* – (*N* + 1)/2) where the scalar *a* ensured the norm of the vector was 1. All columns were made orthogonal using the Gram-Schmidt procedure with *U*_*m*_ columns orthogonalized in the order *m*, 1, 2… *m-1* to preserve most of the structure of columns *m* and 1.

For the connections between Schur directions, we used

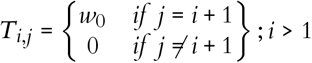

so that the *ith* direction connects only to the *i+1th* direction. The strength of the feedforward connections, *w*_*o*_, was set to 1.06 to produce amplification matched to the data. Additional feedforward connections from the first column *U*_*1*_ were added for the FF only and AFF networks to reduce the dimensionality of the trial-averaged responses and increase the influence of the first direction, to resemble one-dimensional selectivity in ALM (Fig. 1m; Supp. Fig. 1). For the FF only network *U*_*1*_ projected to every 5^th^ vector starting with *U*_*2*_ (i.e. *U*_*7*_, *U*_*12*_, etc) with strength *w*_0_/20. For the AFF network *U*_*1*_ projected to 15 randomly selected directions with strength *w*_0_/10. For the FF only network, τ = 25ms. For the FB only and AFF models τ = 50ms. For the AFF network *T*_1,1_ = 0.9 resulting in a decay of 500 ms along *U*_1_. For the FB only network *T*_1,1_ = 1 resulting in perfect integration along *U*_*1*_.

Robustness of model networks following broad photoinhibition was tested in an AFF model (Supp. Fig. 5) and a non-linear one-dimensional attractor network. The one-dimensional attractor network consisted of two neurons with the connectivity matrix

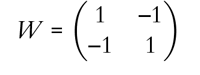

and synaptic non-linearity *f*(*x*) = 5 (tan *h*(*x*/10) + 1). Robustness was tested with 15 different photoinhibition strengths. Each photoinhibition strength was simulated 100 times in the presence of noise to assess the strength of the correlation between CD activity at the end of the delay and during the early delay epoch (−2.0 s < *t < -1*.*4 s*).

The RNN in Supp. Fig. 4 was trained using the FORCE algorithm (Sussillo & Abbott 2009) to generate a ramping output along a single output direction. The network contained 400 units, had τ = 200 *ms* and neuronal activation *f* (*x*) = *tanh*(*x*). The connectivity was randomly initialized with mean zero, connection probability *p* = 0.1 and variance 1.5^2^/(*pN*). In the trained RNN, we identified the CD as the direction that maximized choice selectivity. To identify the behavior influencing subspace we perturbed 500 random directions and characterized the influence that each perturbation had on the CD. We then estimated the optimally influential direction using linear regression to relate the perturbations to the influence. This process was repeated iteratively. When selecting the 500 random perturbation directions during each iteration we required that these directions had zero overlap with the previously identified optimal directions. This method allowed us to identify as many independently influential directions as possible in the network.

### Two-photon imaging and targeted photostimulation data

Two-photon imaging data (Daie et al. 2021a), as described previously (Daie et al. 2021b), were acquired during 84 behavioral sessions from eight mice with an average of 193 imaged neurons (range, 131—275) (Figs. 3-4; Supp. Fig. 3). In these experiments, groups of eight neurons were photostimulated during the delay epoch. Neurons were photostimulated sequentially for 3 ms each with 1 ms gaps between neurons. The cycle of eight neurons was repeated ten times resulting in a 319 ms duration photostimulation. Photostimuli were delivered on 30-40% of all trials with 2-5 different groups per experimental session. The delay epoch was three seconds in these experiments.

Given the spatial resolution of targeted photostimulation, we defined any neuron within 20 μm of a photostimulation target as ‘directly photostimulated’ and all neurons 30 μm or farther from a photostimulation target as ‘coupled’. For each neuron, *i*, and photostimulation group, *pg*, we calculated 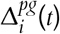 as the difference in activity between photostimulation and non-photostimulation trials. We separately analyzed 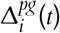 for directlyphotostimulated (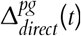, Fig. 4a,d) and coupled (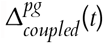, Fig. 4b,c,e,f) neurons.

In calcium imaging data, we identified five directions in activity space: the *TD*_*t*=*s*_, *TD*_*t*=–3*s*_,*TD*_*t*=–2*s*_, *TD*_*t*=–1*s*_ and CD. In the two-photon imaging data set the CD was calculated as the average selectivity in the 1s window centered on the delay end (Daie et al. 2021b). Fewer *TD*_*t*_ were calculated in the imaging data than in the electrophysiological recordings because the temporal resolution in calcium imaging is lower than in electrophysiological recordings. The CD was computed using data from a 1s window centered on the end of the delay. The *TD*_*t*=*s*_ was calculated using data from the sample epoch. *TD*_*t*=–3*s*_, *TD*_*t*=–2*s*_, and *TD*_*t*=–1*s*_ were calculated using 1s windows starting at -3s, -2s and -1s relative to the end of the delay epoch.

### Behavioral bias following targeted photostimulation

For each photostimulation group, we calculated the percentage change in behavioral choice 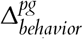 as the change in fraction of rightward choices between photostimulation and non-photostimulation trials (Fig. 3). Next, we calculated the selectivity of all directly photostimulated neurons along the 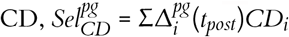 (Fig. 3c) and 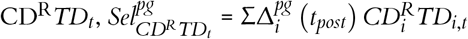. In Fig. 3f the x-axis was 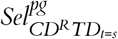. Next, we calculated the correlation between 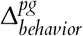 and 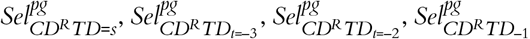 and 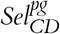 (Fig. 3i).

### Causal connectivity

To calculate the strength of connections from *TD*_*t*_ onto TD_*t’*_ we first identified experiments in which photostimulation strongly activated *TD*_*t*_ and then calculated the changes in activity along TD_*t’*_ in coupled neurons (Fig. 4).

To identify photostimulation groups with strong positive activation along 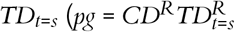; Fig. 4a-c; 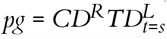, Fig. 4d-f) we restricted our analysis to neurons with positive (rightward) CD selectivity. This choice was based on our previous work showing that neurons with rightward, but not leftward, CD selectivity were strongly coupled to the surrounding network (Daie et al. 2021b). Thus, we define the directions 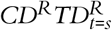 and 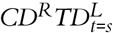.

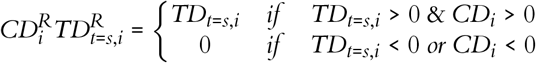

and

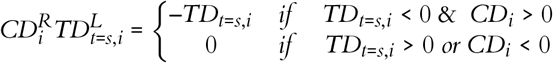

Using these definitions, we calculated the overlap of the photostimulation response of the directly photostimulated neurons and the directions 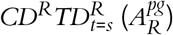 and 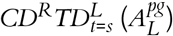 as

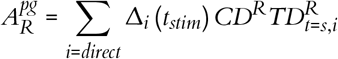

and

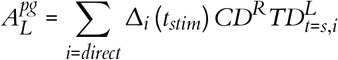

for each photostimulation group where *t*_*stim*_was the 200 ms following the offset of photostimulation. For the plots in Fig. 4a-c we included the photostimulation groups that most strongly activated 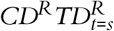 (referred toas 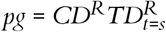) such that 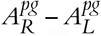 was in the 70^th^percentile across all experiments (mean,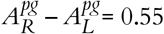, 64 photostimulation groups). For the plots in Fig. 4d-f we included the groups with strong activation of 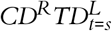 neurons such that 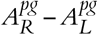 was in the bottom 30^th^ percentile across all experiments (mean 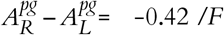, 64 photostimulation groups).

The causal connection matrix 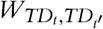 prepresents the strength of the causal connection from direction TD_*i*_ to direction TD_*j*_. For example, the connection strength from CD^*R*^*TD*_*t=s*_ to the 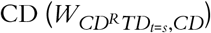 was calculated as the response of the coupled neurons projected along the CD following photostimulation of 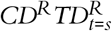 neurons minus their response following photostimulation of 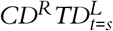 neurons as follows

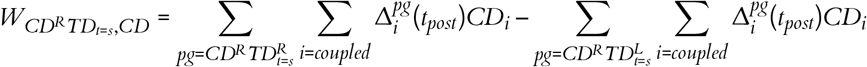

where *t*_*post*_ is the time from 2.0s – 2.9s following the photostimulation. The sums over 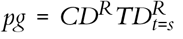 and 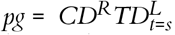 correspond to photostimulation groups in which the *TD*_*t*=*s*_ was strongly activated in the right (positive) or left (negative) direction, respectively.

To assess the spatial structure of the connectivity, we calculated the sums for 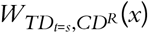 and *W*_*CD,CD*_(*x*) in different spatial bins (Fig. 4h-i). For example

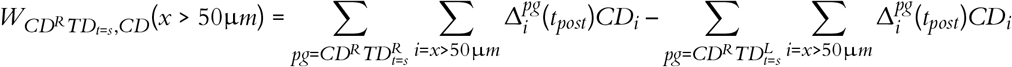

The degree of feedforward bias in the causal connectivity matrix was tested by calculating the Pearson correlation between 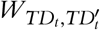 and *t* – *t*’. A positive correlation in this relationship would reflect a feedforward bias where early to late connections are stronger than late to early connections.

## Code availability

Code is available in the Github repository (https://github.com/kpdaie/ALM_feedforward).

## Acknowledgments

We thank S. Romani, H. Inagaki, N. Li, M. Rozsa, J. Cohen and J. Siegle for comments on the manuscript; R. Darshan and T. Wang for helpful discussions. This work was funded by the Howard Hughes Medical Institute (K.D., L.F. and K.S.), the Paul G. Allen Foundation (K.D. and K.S.), the Simons Collaboration on the Global Brain (K.S. and S.D.), the NIH (5U19NS123714 K.S. and EB028171 S.D.), McKnight foundation (S.D.), the Sloan foundation (S.D.), and Excellence Initiative of Aix-Marseille University -A*MIDEX (L.F.).

## Author contribution matrix

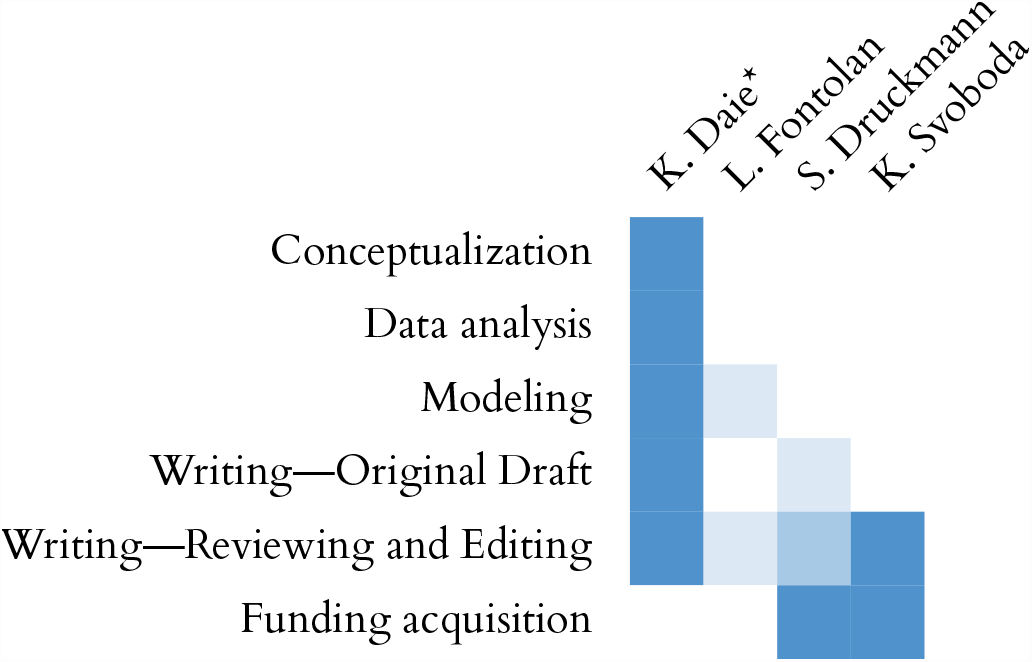

---

**Figure S1:**
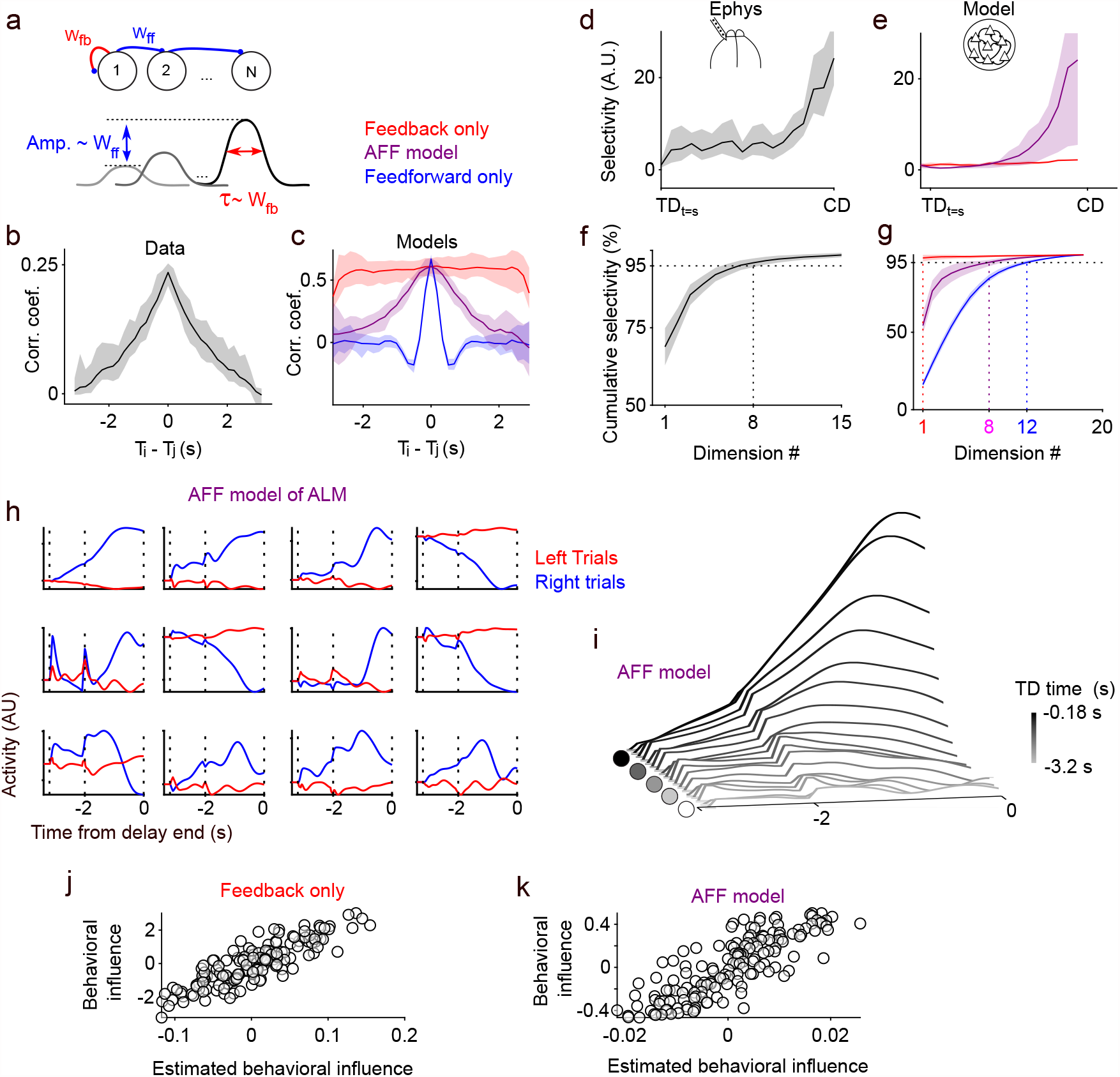
Amplifying Feedforward dynamics in ALM. **a**. Schematic connectivity in an amplifying feedforward network. Feedforward connections contribute to amplification of signals (red) as it propagates down the chain. Feedback connections elongate the response along each direction (blue). **b-c**. Correlation between *TD*_*t*_ and *TD*_*t′*_ vs *t – t’* for data (b) and models (c) with FB only (red), AFF (purple) and FF only connections (blue). Error shade, 95% bootstrap confidence intervals. **d-e**. Variance along each *TD*_*t*_ normalized by variance along the first *TD*_*t*_ for data (e) and AFF model (purple) and FB only model (red). Error shade, 95% bootstrap confidence intervals. **f-g**. Cumulative variance along singular vectors computed using the set of *TD*_*t*_ for data (g) and models. Error shade, s.e.m across experimental sessions. **h**. Responses of individual neurons in the AFF model on lick left (red) and lick right (blue) trials. **i**. Responses along each *TD*_*t*_ for the AFF model. **j-k**. Behavioral influence vs. estimated behavioral influence (*TD*_*t*_) for a feedback only model (j) and the AFF model (k). All data were averaged across 19 recording sessions from 6 mice. Models were average across 20 simulations.

**Figure S2:**
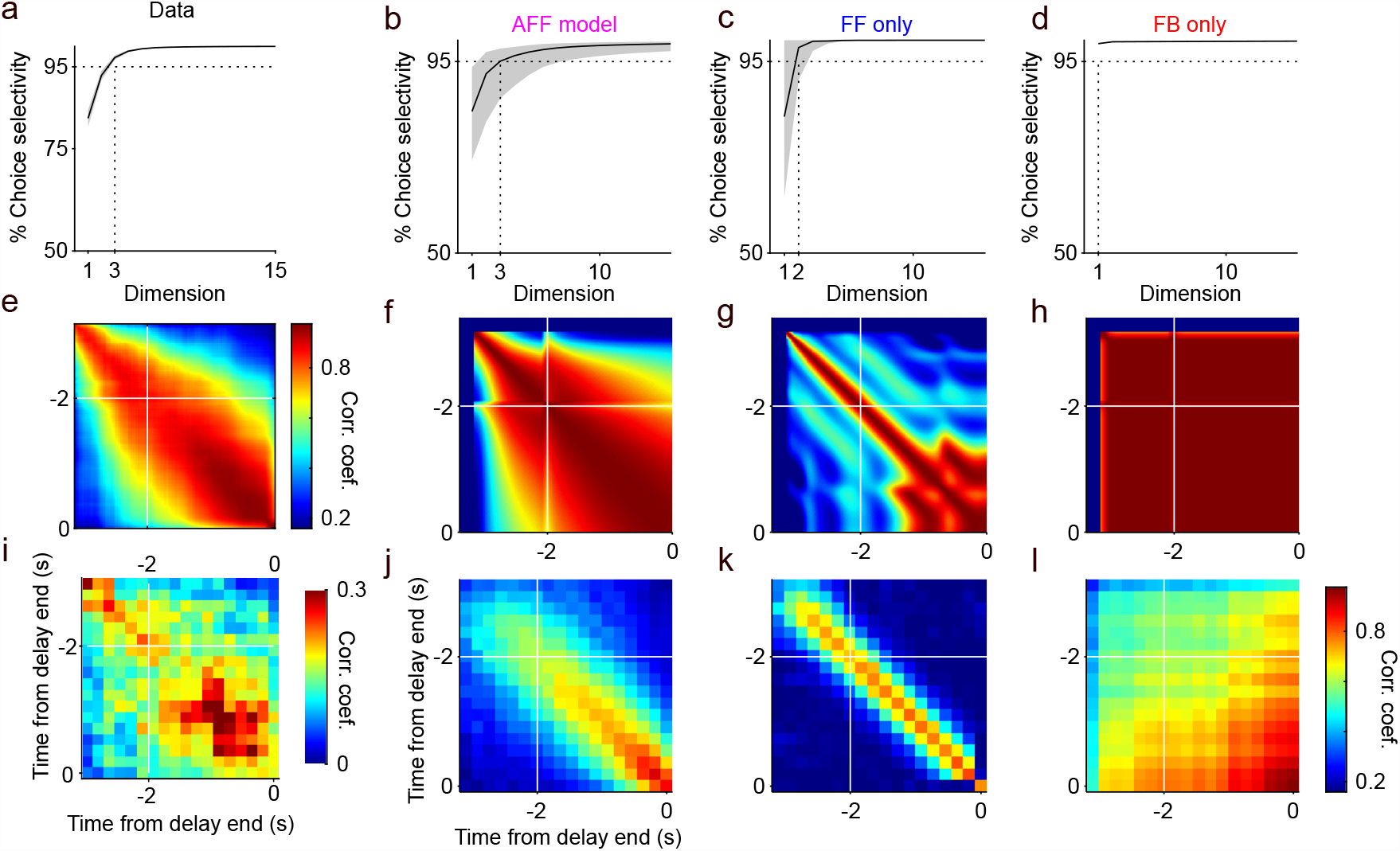
Dimensionality of ALM data and model dynamics. **a-d**. Dimensionality of trial-averaged choice selectivity (difference in activity on Right and Left trials) for ALM data (a) and model networks (b-d). Error shade, 95% bootstrap confidence intervals. **e-h**. Correlation between trial-avg. selectivity at time point *t* vs time point *t’* for all sample and delay epoch time points. **i-l**. Correlation *TD*_*t*_ and *TD*_*t′*_ for all sample and delay epoch time points.

**Figure S3:**
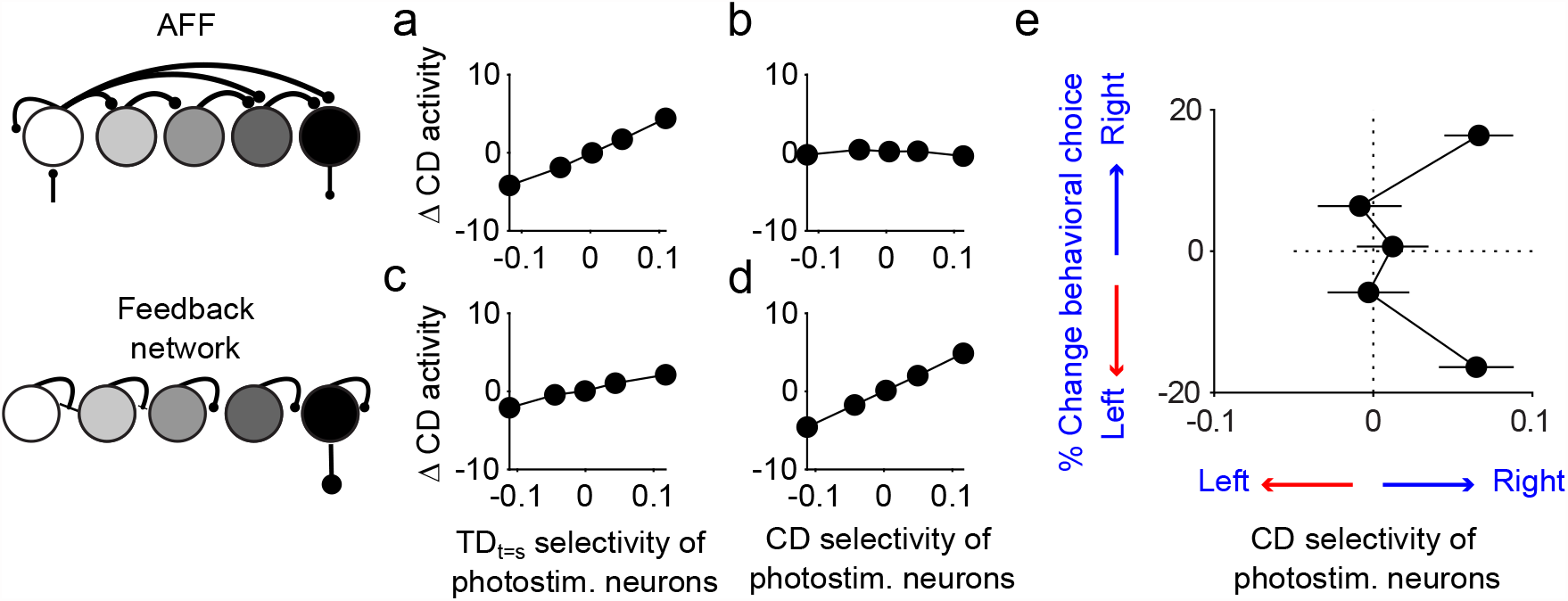
Behavioral bias following photostimulation. Response of AFF (a-b) and FB only (c-d) model networks to brief photostimulation of 2 neurons. **a-d**. Photostimulation-induced change in activity along CD (Δ CD activity) vs. overlap of photostimulation with *TD*_*t*=*s*_ (a,c) and CD (b,d). Δ CD activity was correlated with photostim. *TD*_*t*=*s*_ overlap for both the AFF (Pearson correlation = 0.8; p = 2×10-23) and FB only network (Pearson correlation = 0.56; p =1.6×10-9). Δ CD activity was correlated with photostim. CD overlap for FB only network (Pearson correlation = 0.98; p = 0) but not for the AFF (Pearson correlation = -0.02; p = 0.89). Note that *TD*_*t*=*s*_ and CD are nearly identical for the FB only model with noise (Fig. 1f) and exactly identical without simulation noise (not shown). **e**. Change in behavioral choice vs. selectivity of directly photostimulated neurons along CD. Same data as Fig. 3c except data were plotted in 5 equally spaced bins along the y-axis. Error bars, s.e.m across photostimulation groups.

**Figure S4:**
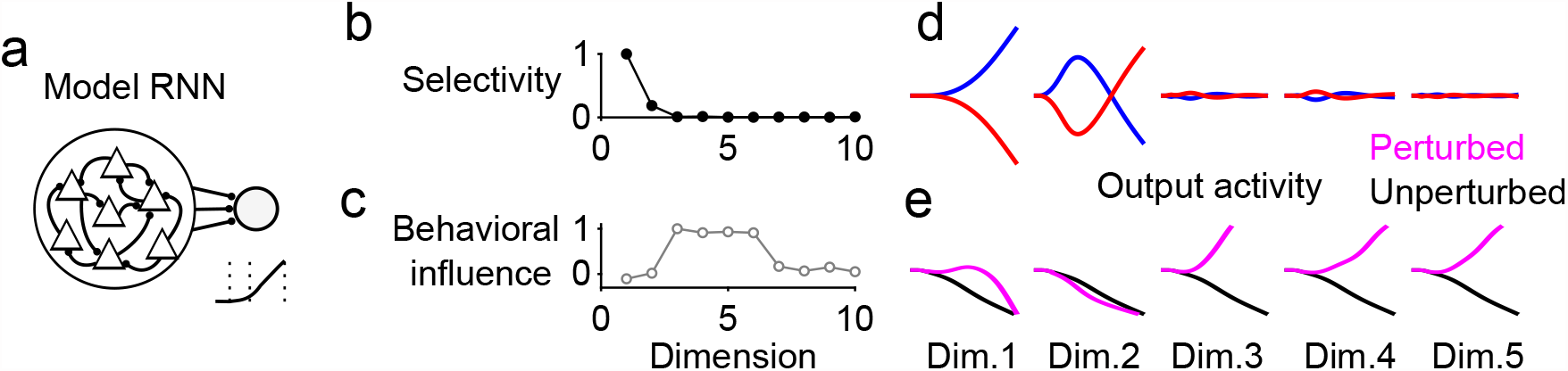
Figure 4: RNN model of ALM. **a**. RNN schematic, trained to produce ramping output. **b**. Amplitude of selectivity within a ten-dimensional subspace. First two dimensions are the first two dimensions from a singular value decomposition. Dimensions 3-10 were identified as having the strongest influence on behavior (Methods). **c**. Behavioral influence of the same 10 dimensions as in panel b. **d**. Activity projected along the first five dimensions. **e**. Activity along the CD when one of the first five dimensions was perturbed (magenta) compared to CD activity without perturbation (black).

**Figure S5:**
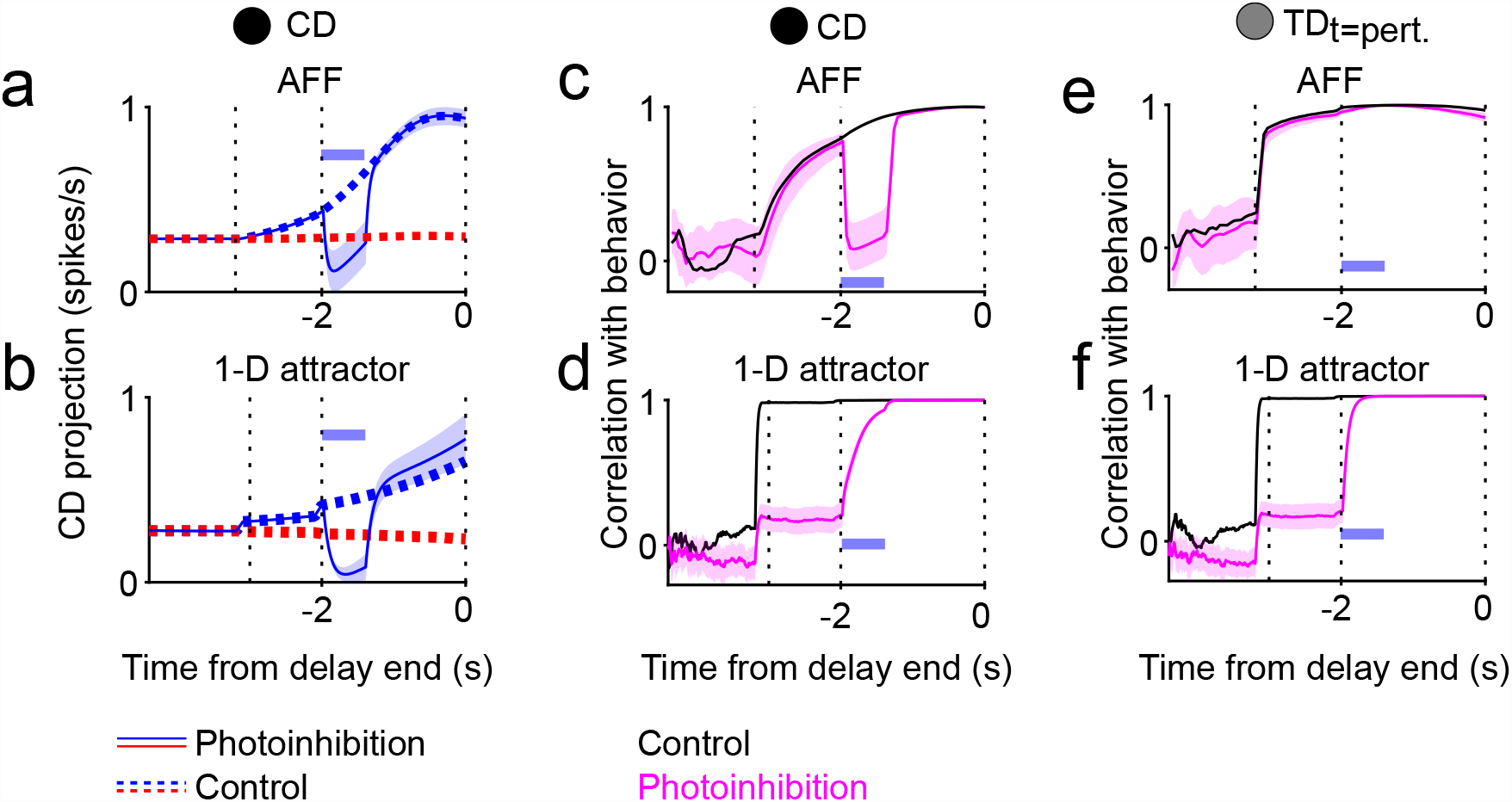
Activity along behavior influencing subspace during perturbations in network models. **a-b**. Activity along CD on control (dashed), lick right (blue), lick left (red) trials and lick right photoinhibition trials (solid) an AFF model (a), and a 1-D attractor model (b). **c-f**. Correlation between activity projections along CD (c-d) and *TD*_*t*=*pert*._ (e-f) an AFF model (c,e) and a 1-D attractor model (d,f) during control (black) and photoinhibition (magenta) trials. Blue bars, time of photoinhibition.

